# SOX10 and microRNAs: decoding their interplay in regulating melanoma plasticity

**DOI:** 10.1101/2024.10.09.617381

**Authors:** Xin Lai, Chunyan Luan, Zhesi Zhang, Anja Wessely, Markus V. Heppt, Carola Berking, Julio Vera

## Abstract

Recent studies show that the dysregulation of the transcription factor SOX10 is essential for development and progression of melanoma. MicroRNAs (miRNAs) can regulate the expression of transcription factors at the post-transcriptional level. The interactions between SOX10 and its targeting miRNAs form network motifs such as feedforward and feedback loops. Such motifs can result in non-linear dynamics in gene expression levels, therefore playing a crucial role in regulating tumor proliferation and metastasis as well as tumor’s responses to therapies. Here, we reviewed and discussed the intricate interplay between SOX10 and miRNAs in melanoma biology including melanogenesis, phenotype switch, and therapy resistance. Additionally, we investigated the gene regulatory interactions in melanoma, identifying crucial network motifs that involve both SOX10 and miRNAs. We also analyzed the expression levels of the components within these motifs. From a control theory perspective, we explained how these dynamics are linked to the phenotypic plasticity of melanoma cells. In summary, we underscored the importance of employing a data-driven network biology approach to elucidate the complex regulatory mechanisms and identify driver network motifs within the melanoma network. This methodology facilitates a deeper understanding of the regulation of SOX10 by miRNAs in melanoma. The insight gained could potentially contribute to the development of miRNA-based treatments for SOX10, thereby enhancing the clinical management of this malignancy.

## 1. Introduction

### 1.1 Cutaneous Melanoma

Cutaneous melanoma is an aggressive and deadly skin cancer, resulting in 57,000 deaths in 2020 (1). In the last decade, its incidence has steadily increased, especially in Caucasian populations, imposing great challenges for global healthcare (2, 3). Clinical phenotype, genetic background, and UV exposure play a significant role in affecting geographic variations in incidence and mortality of melanoma (4). Currently, it is one of the most common cancers in fair-skinned people, especially those with blond or red hair and light-colored eyes. Unlike other solid tumors, melanoma is one of the most common types of cancer diagnosed in young adults (5–7). The long-term survival rate for patients with metastatic melanoma was only 5% (8) and has improved significantly since the introduction of BRAF/MEK inhibitors and immune checkpoint blockade after 2010 (9). There is a strong correlation between survival rates and the stage of melanoma at the time of diagnosis (10). The patient’s prognosis is also affected by factors such as genetic heterogeneity of the tumor and patients’ access to advanced therapeutic approaches. For example, systemic therapeutic approaches such as small molecule inhibitors of mutant BRAF kinase and its downstream target MEK, but especially immune checkpoint inhibitors, have drastically changed the landscape of melanoma therapy over the past decade (11). These treatment options significantly improve survival rates for patients with metastatic melanoma (12–14). Currently, 5-year survival rates of 30-50% can be achieved in clinical trials in advanced and metastatic melanoma (9). However, the mortality of melanoma patients remains high and the incidence of metastatic melanoma is growing (15–20). Therefore, advanced techniques have been used to develop complex melanoma models to improve understanding of the tumor and facilitate the development of therapeutics (18).

Recent research into gene regulatory networks of tumor has provoked a great interest in the oncology community (21, 22). The underlying idea is that cancer driver genes do not act in isolation but are integrated into large and interconnected signaling and transcriptional networks. In fact, the identification of novel genes from molecular networks, and targeting them using pharmacological drugs or genetic intervention methods are considered the next major step forward in the field of cancer therapy (23–26). Because these networks are large and contain many interconnected complex gene circuits, the potential targets for cancer could be derived from network biology analysis, a family of data-and systems-driven computational methods used to characterize the molecular interactions and their dynamics underlying the tumor. Here, we showcased the advance of network biology methods in analyzing the interactions between SOX10 and miRNAs in melanoma and provide quantitative elaboration on the regulation of SOX10 in miRNA-mediated network motifs.

### 1.2 SOX10

SOX10 belongs to the HMG-box family of transcription factors (TFs) and plays an essential role in the development of melanocytes and other neural crest (NC)-derived cells (27, 28). SOX10 is expressed in the very early phase of NC formation before the migration of NC cells starts, and its expression is maintained throughout the whole migratory phase of melanocytic precursors (27, 29, 30). SOX10 is critical for the transition of pre-migratory cells to migrating NC cells and is essential for NC cell survival (31). Specifically, the loss of SOX10 significantly inhibited the generation of post-migratory NC stem cells, increased the cell apoptosis rate during NC commitment, and severely compromised both neuronal and glial differentiation capacities. Moreover, the multipotent post-migratory NC cells that form and migrate in the absence of SOX10 undergo apoptosis before reaching maturation stage, implicating SOX10 as an essential gene in regulating the survival of undifferentiated NC cells (28). In addition to its function in maintaining NC formation, SOX10 is expressed in NC cells to convert them to the melanocytic lineage by directly upregulating the expression of MITF (32–35).

Melanoma is a malignant transformation of melanocytes and becomes more invasive and resistant to available treatments after gaining metastatic ability (36). Common driver mutations, such as the BRAF V600E mutation, are prevalent in melanoma (37, 38). These mutations lead to constitutive activation of the MEK-ERK signaling cascade. However, it’s important to note that BRAF mutations alone are not sufficient to induce melanoma due to a phenomenon known as oncogene-induced senescence and that BRAF mutations are also often found in many benign nevi. The exact course of events in which melanocytes become tumorigenic remains unclear. The concept of cancer stem cells is coined by Nishimura et al. (39) and provides a potential sound explanation for melanoma onset. In particular, during melanoma formation, melanocytes acquire stemness phenotype (40) characterized by the expression of stem cell markers such as NES (41), NGFR (42), and PROM1 (43). Melanocytes can reacquire NC characteristics, become stem-like and less differentiated, which promotes cell proliferation, migration, and metabolic changes (44). The expression of the stem cell markers is usually accompanied by metabolic changes in melanocytes (45). For example, in the process of melanomagenesis and in the context of oxygen deprivation in tumors, melanoma cells often switch their energy production from mitochondrial metabolism to glycolysis because the by-products of glycolysis can be conveniently redirected to critical pathways for the synthesis of macromolecules (46). The progression of melanoma and its resistance to treatment is also associated with dysregulated sphingolipid metabolism (47). Due to the genetic and metabolic alternative, melanocytes undergo multiple divisions in an asymmetrical or symmetrical manner and acquire the phenotype of uncontrolled proliferation. For instance, loss of CDKN2A, TP53, and PTEN expression are often found in melanoma, leading to uncontrolled cell division (44). Furthermore, it is rational to assume that melanocytes with stemness phenotypes are the seeding tumor cells and can further develop into primary tumor (48). The acquisition of the stemness phenotype in melanocytes was found to be potentially dependent on the expression of SOX10, which can regulate the expression of the aforementioned stem cell markers (42, 43, 49, 50). Taken together, this suggests that SOX10 is a crucial factor for melanocytes to acquire stemness phenotype and their development into tumor cells.

In melanoma, SOX10 plays a diverse role related to the proliferative, metastatic, and immunogenic capacity of tumor cells. Specifically, SOX10 is critical for the proliferation of melanoma cells. It has been observed that a decrease in SOX10 levels not only reduces proliferation but also increases the invasive properties of these cells (51). In addition, SOX10 plays a critical role in the migration of melanoma cells. This was demonstrated by a significant reduction in migration activity when melanoma cells were transfected with siRNAs specific for SOX10 (52). In addition to its role in cell migration, SOX10 has been implicated in cell cycle regulation in melanoma. Tumor cells with stable loss of this gene were found to be arrested in the G1 phase of the cell cycle (53). In addition, SOX10 has been identified as a regulator of the expression of immune checkpoint proteins such as HVEM and CEACAM1 in mice (54). It also has a role in modulating melanoma immunogenicity through an IRF4-IRF1 axis (55).

Moreover, SOX10, in conjunction with other NC factors, forms complex regulatory networks in melanoma. The SOX10-MITF pathway plays a crucial role in melanoma cell survival, proliferation, and metastasis formation (56). The undifferentiated melanoma subtype that does not show typical melanocytic markers (e.g. SOX10) is characterized by the loss of the melanocytic program, which is primarily driven by the SOX10-MITF axis (133, 134). Additionally, MITF is regulated by PAX3 and TFE3, and is essential for melanogenesis, cell cycle progression, and melanoma cell survival (57, 58). PAX3 plays a role in early NC cell development and influences SOX10 and MITF expression (59). Aberrant PAX3 expression is associated with melanoma advancement and therapy resistance (60). TFE3 is implicated in melanocyte development and pigmentation, with its gene fusions contributing to specific melanoma subtypes (61). In addition, other NC factors, such as SNAIL and SLUG, are involved in the epithelial-to-mesenchymal transition, which promotes melanoma cell invasiveness and metastasis (58, 59).

Hence, expanding our knowledge about the function of SOX10 in melanoma may facilitate the development of novel therapies. It could be particularly helpful in reducing melanoma cells’ stemness properties that endow them with abnormal self-renewing ability and drive malignant transformation (42, 43, 49, 50, 64–66).

### 1.3 MicroRNAs

MicroRNAs (miRNAs) are short, endogenous, non-coding RNAs (ncRNAs) with the ability to bind to 3′-untranslated region (3′-UTR) of target messenger RNAs (mRNAs), and can repress their target gene expression by affecting the stability of mRNAs or inhibiting protein translation (67–70). Some studies show that miRNAs can repress the transcription of target genes by binding to 5′-untranslated region (5′-UTR) of mRNAs (71–74). It has also been shown that miRNAs can transcriptionally modulate gene expression through *de novo* DNA methylation (75). Due to the ability to regulate the expression of vast protein-coding genes, miRNAs have emerged as key players in controlling cell homoeostasis and phenotypes (76–78). Accumulative evidence show that these small ncRNAs are involved in melanoma initiation and progression via different mechanisms (79–82). Furthermore, dysregulated endogenous and/or circulating miRNAs have been linked to the pathogenesis of melanoma, including the process of tumor invasion and metastasis, angiogenesis, the establishment of microenvironment, and immune escape (34, 79–81, 83–86). The pivotal role of miRNAs in regulating genes associated with angiogenesis, stemness, proliferation, apoptosis, and resistance to treatment (87–90) in melanoma suggests they have potential for clinical applications as prognosis and diagnosis biomarkers, as well as therapeutic targets (91–93).

miRNAs can regulate multiple targets involved in malignancy, either by controlling a single gene target (94) or multiple ones simultaneously (24, 25, 27). Due to the promiscuous gene targeting, miRNAs can regulate gene expression in a cooperative manner (95). It has been shown that cooperative miRNAs can synergistically repress the expression of their target genes to regulate tumor phenotypes such as chemoresistance in melanoma cells (96, 97). Also, key cancer genes like CDKN1A are regulated by multiple miRNAs alone or in a concerted manner (98). In melanocytes, multiple miRNAs can collectively target one particular gene, such as MITF that is a transcriptional target of SOX10, for fine-tuning its expression (99–102), therefore regulating signaling pathways associated with cell proliferation and migration (103, 104). In this review, we focused on the regulation of SOX10 by miRNAs in melanoma. To characterize the role of such interactions in melanoma, we elaborated on how they can affect the growth, proliferation, and migration of tumor cells.

## 2. The role of miRNAs and SOX10 in melanoma cell biology

The TF SOX10 can induce stemness characteristics in melanoma cells and sustains their proliferative and tumorigenic capabilities. However, no comprehensive review has been conducted to elucidate the interactions between SOX10 and miRNAs in the context of melanoma. Here, we provided a detailed analysis of the most relevant studies that have explored the role of SOX10, miRNAs, and their interacting molecules such as MITF that plays a multifaceted role in melanoma includes regulating gene expression, influencing tumor heterogeneity, affecting immune cell attraction, and presenting potential therapeutic targets (105–107) (**Figure 1**). Since these genes have significant implications for melanoma, it is worth discussing their impact on the phenotypic attributes of melanoma cells. Furthermore, by collecting facts from published reports, we reconstructed the state-of-the-art on the molecular interplay between SOX10, miRNAs, and other interacting molecules is thoroughly examined in melanoma.

**Figure 1.**
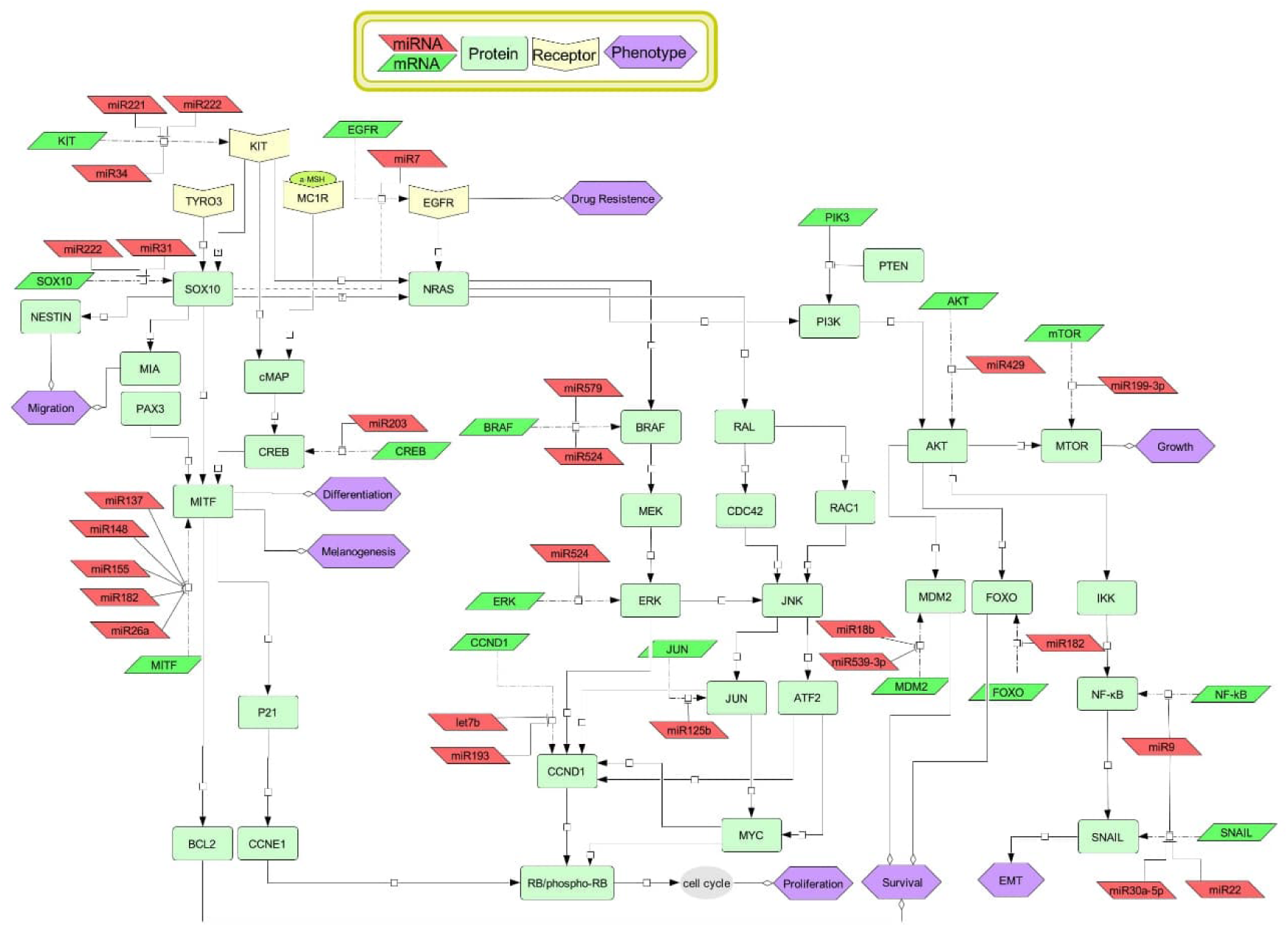
Overview of the role of SOX10 and miRNAs in melanoma. The signaling pathways and gene regulatory interactions related with melanoma were extracted from the literature and are described and discussed in the main text. The figure was created in CellDesigner, a software tool that employs a systems biology graphical notation to facilitate the exchange of graphical depictions of molecular pathways. The nodes account for molecular species and triggered phenotypes (see the figure legend). A CellDesigner file in xml format is provided in supplementary materials and the references used for annotating the molecular interactions are accessible in Table S1. EMT: epithelial-mesenchymal transition.

### 2.1 Melanogenesis

Melanoma develops from melanocytes (108), cells that contain specialized organelles called melanosomes. These organelles contain two types of pigments: pheomelanin and eumelanin (109). The production of eumelanin, which is the primary photoprotective mediator, is promoted by the binding of α-MSH to MC1R. This phenomenon leads to a significant increase in intracellular cAMP levels and the activation of CREB (110–112). Together, SOX10 and PAX3 induce the transcription of MITF, which is commonly known as the master regulator of melanogenesis, a multi-enzymatic process that starts with the amino acid L-tyrosine and ends with melanin synthesis (113, 114). MITF plays a direct role in promoting the transcription of genes encoding for enzymes that are essential for melanogenesis, such as TYR and DCT (114).

MiRNAs play a role in the regulation of melanogenesis and melanocytic neoplasms (115, 116). Numerous studies have investigated miRNAs that affect MITF expression, thereby regulating mRNA expression levels of melanogenic enzymes. For instance, research indicates that the cluster composed of miR-182, miR-183 and miR-96 targets the 3′-UTR of MITF in B16 mouse melanoma cells (117). The overexpression of the miRNA cluster resulted in a decrease in the expression of MITF, its transcriptional target genes TYR, TYRP-1, and DCT, as well as in the reduction of melanin production. Conversely, the knockdown of the miR-183 cluster increased the expression of MITF, TYR, TYRP-1, and DCT, leading to an increase in melanin levels. Similar effects were observed when knocking down miR-143-5p, which resulted in increased expression of the above mentioned melanogenesis genes with MLPH and RAB27A (118). Other miRNAs, such as miR-340, miR-141-3p, and miR-200a-3p, have also been studied for their role in regulating MITF expression and melanogenesis (117). In addition to these miRNAs, which play significant regulatory roles in MITF regulation, other miRNAs are involved in melanogenesis by regulating directly the expression of melanogenic enzymes, as well as other TFs or components of the signaling pathways that regulate melanogenesis (116). For example, miR-203 targets KIF5B to influence melanocyte pigmentation (119) and miR-211 targets TGFBR2, which modulates melanogenesis by affecting tyrosinase and TYRP1 (120).

Several miRNAs have been identified to target SOX family members. For instance, miR-21a-5p overexpression downregulated SOX5 in normal human melanocytes (121). miR-140-5p, found to be downregulated in melanoma tissues and cells, directly targets SOX4 and inhibits its expression, thereby hindering melanoma cell proliferation and invasion (122). Changes in miR-145 expression are positively correlated with SOX9 expression, a SOX family member involved in melanin biosynthesis (123). Another miRNA that targets SOX family members is miR-200c that targets SOX1. Exosomal miR-200 from keratinocytes downregulates SOX1 resulting in increased expression level of β-catenin in the nucleus that upregulates the transcription of MITF and its target melanogenesis genes TYR, TRP1 and TRP2 (124). miR-155 can repress the expression of melanogenesis associated genes including SOX10 in melanocytes and keratinocytes (125). SOX10 plays a role in regulating the SOX10-MITF pathway, which contributes to melanogenesis by regulating melanoma cell survival, proliferation, and metastasis formation (56). Therefore, further investigation is required to better understand the role of miRNAs and SOX10 in regulating melanocyte development and pigmentation.

### 2.2 Melanoma cell proliferation, growth, and metastasis

Numerous studies have demonstrated the significant role of SOX10 in melanoma growth. Human melanoma cells with stable SOX10 knockdown exhibited arrested cell growth, altered cell morphology, and induced senescence (126). Additionally, SOX10 can promote the formation and maintenance of giant congenital naevi and melanoma (64). There are also studies that have confirmed the role of SOX10 in inhibiting melanoma cell proliferation through the Notch signaling pathway (127) or the WNT/β-catenin signaling pathway (128). A study showed that the receptor tyrosine kinase KIT is upregulated in melanoma (129). KIT is a crucial mitogen and survival factor for melanocytes and melanoblasts. Although activating mutations in the KIT gene are infrequent, their oncogenic properties may play a significant role in melanoma development when they occur (64, 130–133). Rönnstrand and Phund observed disparate expression patterns of SOX10 and KIT during melanoma progression. SOX10 expression levels exhibited a gradual increase throughout the progression of melanoma, whereas KIT expression was upregulated in primary melanoma but reduced in metastatic melanoma (133). Shakhova et al. showed that upregulation of SOX10 is associated with expression of the NRAS oncogene in mice and SOX10 haploinsufficiency impeded the proliferation of tumor cells with overexpressed NRAS (64, 65). Consequently, this could represent an efficacious approach to halting melanomagenesis in genetically predisposed individuals who are at an elevated risk of developing this tumor due to the presence of the NRAS mutation. Additionally, SOX10 was found to play a role in the activation of the MAPK and PI3K/AKT signaling pathways that are known to regulate cell growth, proliferation, and survival in melanoma (134, 135). I*n vitro* and *in vivo* invasion assays suggest the involvement of SOX10 in melanoma cell spreading (136). The results showed a significant reduction in the invasion capacity of melanoma cells when SOX10 was inhibited by small inference RNA. This indicates that SOX10 plays a crucial role in the invasion of melanoma cells, possibly by affecting the expression of NES (137, 138) and MIA (139) whose expression are correlated with melanoma progression.

There is increasing evidence that miRNAs can act as tumor suppressors to regulate the proliferation and growth of melanoma (140), as well as melanoma metastasis (141). A recent study investigated the relationship between miR-31 and SOX10 in melanoma growth. Upregulation of SOX10 and downregulation of miR-31 were found in primary melanoma tissues and SOX10 was further identified as a target of miR-31 (142). In the experiments, overexpression of SOX10 dramatically promoted melanoma cell proliferation, while increased expression of miR-31 suppressed cell growth. Investigation of the molecular pathways behind the interplay between SOX10 and miR-31 suggested that the tumor suppressor role of miR-31 was achieved through repression of SOX10 and subsequent downregulation of the PI3K/AKT signaling pathway. A study identified a mechanism involving the MIR222/SOX10/NOTCH axis, where curcumin increases miR-222-3p levels to reduce SOX10 expression, resulting in inhibiting the growth and metastasis of melanoma (143). The results indicate that miR-222 plays a role in regulating the Notch signaling pathway, which is associated with the progression of primary melanoma. This is achieved by activating the MAPK and PI3K/AKT pathways and increasing N-cadherin expression (144). However, the Notch pathway may also have the potential to suppress melanoma with the BRAF V600E mutation and loss of PTEN (144). It showed that miR-107 can regulate melanoma invasiveness (145). The authors found that this miRNA was significantly downregulated in melanoma and had the lowest expression in the metastatic form of the tumor. They showed that exogenous overexpression of miR-107 reduced melanoma cell proliferation, migration, and invasion and that POU3F2 was a downstream target of miR-107 mediating is effect in invasion of melanoma. Interestingly, phosphorylated POU3F2 can control melanocyte proliferation and migration through its interaction with SOX10 and MITF (146). Thus, miR-107 could affect MITF and SOX10 invasiveness activity by modulating the expression of POU3F2. Similarly, miR-211 is consistently downregulated in melanoma cells compared to melanocytes and suppresses melanoma growth and migration via repression of KCNMA1 and MITF (147). miR-211 also inhibits epithelial-mesenchymal transition (EMT) in melanoma cells through repressing RAB22A (148). The MIR222/MIR221 cluster appears as a key regulator in melanoma. Upon activation of the Notch signaling pathway in melanoma cells, MITF de-represses the miRNA cluster, leading to tumor invasion through a mechanism involving the inhibition of GRB10 or ESR1 (149). In conclusion, multiple miRNAs, alone or in combination, have significant implications in melanoma metastasis and invasion due to their ability to regulate the expression of SOX10, MITF and their mutual targets directly or indirectly.

### 2.3 Phenotype switch in melanoma

Melanoma cells are known for their high plasticity and ability to transition between different states, allowing them to adapt to suboptimal conditions and to resist therapies (150–153). This adaptability is demonstrated through the switching between two primary states: the proliferative/differentiated phenotype, also known as the melanocyte-like state, and the invasive/dedifferentiated phenotype, referred to as the mesenchymal-like state (154). The melanocyte-like phenotype is characterized by the expression of melanocytic lineage TFs such as MITF, SOX10, ZEB2, and downstream markers involved in cell differentiation and pigmentation (TYR, TYRP-1, and Melan-A) (155). In contrast, the mesenchymal-like phenotype is defined by the expression of TEADs, the receptor tyrosine kinases AXL and EGFR, the TFs SOX9, ZEB1, BRN2, and genes involved in the WNT5A and TGF-β pathways implicated in cell invasion (155).

Extensive research has been conducted on the mechanisms that underlie phenotypic plasticity in melanoma cells. Several key regulators of phenotype switch have been identified, including MITF, AXL, ROR2, and SOX10. MITF functions as a central metabolic sensor, integrating cell-extrinsic nutrient signals with cell-intrinsic metabolic responses. The integration described drives melanoma cells towards specific phenotypes that enhance their survival under dynamic microenvironmental conditions (156). MITF is also a key player in melanoma biology controlling many aspects of the melanocytic lineage expression through the interplay between the MITF-MIR211 axis and the PAX3-POU3F2 axis (157). MITF and AXL are markers of heterogeneous melanoma phenotypes (150). High MITF and low AXL expression levels are associated with a proliferative phenotype, while low MITF and high AXL expression levels are associated with an invasive phenotype. The signaling receptor ROR2 has recently been proposed as a significant driver of phenotype switch in melanoma. It has been demonstrated to impair cell proliferation while promoting migration, EMT, and chemoresistance (158). Specifically, the signal triggered by ROR2 has been shown to modulate the activities of several TFs, including AP-1, NFAT, SNAIL, ZEB1, TWIST, and SLUG. Additionally, SOX10 has been shown to mediate phenotypic switching in melanoma by inducing a targeted inhibitor-tolerant state, which is likely a precursor to the acquisition of resistance (51). SOX10-deficient cells in melanoma show invasive characteristics and resistance to BRAF and MEK inhibitors but exhibit sensitivity to cIAP1/2 inhibitors that induce cell death. Also, upon knockdown of SOX10, a phenotypic switching from melanocyte-like cells to mesenchymal-like cells was observed at the single-cell level in nine patient-derived melanoma and A375 cell lines (154). That was due to sequential and recurrent of SOX10-mediated gene regulatory networks.

miRNAs are involved in melanoma plasticity through direct or indirect interactions with those phenotype regulators. For instance, miR-182 promotes an invasive phenotype in melanoma by repressing FOXO3 and MITF, whose high expression levels are required for maintaining the melanocyte-like phenotype in melanoma (102). MITF can mediate the biogenesis of the MIR99A/LET7C/MIR125B cluster by binding to TRIM28, a transcriptional elongation factor, to alter the distribution of RNA polymerase II, resulting in the differential expression of the miRNA cluster. The overexpression of miR-99a and miR-125b in proliferative melanoma cells shifts the tumor cells towards an invasive phenotype, while the overexpression of miR-let-7c in invasive melanoma cells induces a transition to a proliferative state (159). Additionally, the overexpression of miR-410-3p caused by endoplasmic reticulum stress in melanoma favors a phenotypic transition of the tumor to an invasive phenotype (160). In conclusion, miRNAs and SOX10 play pivotal roles in regulating the phenotype switch of melanoma. In most cases, such regulation necessitates the involvement of MITF, underscoring the complex interplay of these factors in melanoma plasticity. However, the interaction between miRNAs and SOX10 to regulate phenotype switch in melanoma has not been documented.

### 2.4 Therapy resistance of melanoma

The last decade has seen a significant advance in melanoma treatment with the development of BRAF inhibitors (BRAFi) that target the MAPK signaling pathway, thereby improving patient survival (161, 162). However, a major clinical challenge is the rapid progression and relapse of melanoma approximately six months after BRAFi treatment (163). This resistance to BRAFi can be attributed to the (re)activation of several oncogenic signaling pathways, including the MAPK/ERK and the PI3K/AKT pathways. These pathways can be altered by direct genetic alterations, upregulation of receptor tyrosine kinases, or alternations in downstream signaling in the context of BRAFi resistance (164). Some results indicate that the silencing of SOX10 and MITF is a key factor in inducing therapeutic resistance in melanoma (165). Melanoma subtypes with low MITF expression are often associated with resistance to targeted therapy (166, 167), and around 50% of recurrent melanoma cases exhibit reduced MITF expression (168). Similarly, decreased SOX10 expression in melanoma has been linked to drug resistance (169). SOX10 is crucial for activating the MAPK and PI3K signaling pathways by regulating the transcription of their components such as FOXD3 (170), and its re-activation can lead to treatment resistance in melanoma due to melanoma cell dedifferentiation (171). This resistance is characterized by decreased sensitivity to drugs like vemurafenib that inhibits the MAPK/ERK pathway, as well as an increase in the quiescence and migratory properties of tumor cells. Studies have identified dysregulated expression levels of EGFR as a potential indicator of melanoma cell response to drugs (172). Acquired EGFR overexpression confers BRAFi resistance in BRAF-mutant melanoma, and the downregulation of SOX10 increased EGFR and PDGFRB expression and vemurafenib resistance (173). In melanoma cells that have developed resistance to vemurafenib, the loss of SOX10 may also contribute to resistance to an immunotherapy called oncolytic viruses (174). These viruses have the potential to reverse cancer-associated immune suppression and to stimulate antitumor immune responses (175). Thus, SOX10 plays a crucial role in mediating both adaptive and acquired resistance of BRAF-mutated melanoma cells by regulating the RAS/RAF/MEK/ERK and PI3K/AKT signaling pathways implicated in drug resistance (172). Furthermore, methylation-mediated repression of MITF and SOX10 can contribute to dedifferentiation of melanoma, and the dedifferentiated tumor cells may explain therapy resistance of melanoma (176). The complex interplay between the transcriptional factors (i.e. SOX10 and MITF) and receptors (i.e. AXL and ERBB3) with their ligands (i.e. NRG1 and GAS6) may form a sophisticated mechanism involved in regulating BRAFi-induced resistance in melanoma (177).

miRNAs are also important in regulating the acquisition of drug resistance in melanoma (178–180). The expression of miR-410-3p is upregulated by the BRAFi vemurafenib in melanoma cells, and its overexpression contribute to the development of resistance of the tumor cells to the drug (160). SOX10 and miR-7 may jointly modulate the response of melanoma cells to BRAFi. miR-7, which is significantly downregulated in vemurafenib-resistant melanoma cells, can reverse BRAFi resistance when its expression is restored (181). Mechanistically, miR-7 targets EGFR, IGF-1R, and CRAF, thereby inhibiting the MAPK and PI3K/AKT signaling pathways. Both SOX10 and miR-7 can regulate EGFR at the transcriptional (173) and post-transcriptional (181) levels, respectively, and EGFR overexpression can lead to BRAFi resistance (182). Therefore, the repression of EGFR by overexpressing both SOX10 and miR-7 could exert synergistic effects on sensitizing the response of melanoma cells to BRAFi. In addition, there are other miRNAs involved in the regulation of drug resistance in melanoma. Some miRNAs exert their effects by targeting MITF, while others directly target genes associated with drug resistance, such as BRAF (183). A recent study demonstrated that the reciprocal interplay between miR-579-3p and MITF in BRAF-mutant melanoma cells. Experiments showed that long-term resistance to the targeted therapy MAPK inhibitor was observed in tumor cells characterized by the loss of both miR-579-3p and MITF (184). This suggests the involvement of the MITF-MIR579 axis in the development of resistance in BRAF-mutant melanoma cells. In addition to SOX10 and MITF, miRNAs may regulate drug resistance in melanoma by targeting other genes. For instance, the downregulation of miR-92a-1-5p has been associated with acquired resistance to BRAFi, as this miRNA inhibits cell viability by targeting S100A9 (185). The downregulation of IGF1R by overexpressed miR-30a-5p has been linked specifically to cisplatin resistance in two melanoma cell lines, demonstrating the influence of the miRNA on chemoresistance in melanoma (186). In conclusion, miRNAs may play significant roles in the development of treatment resistance in melanoma, particularly in relation to SOX10 and MITF.

## 3. Network Motifs

In cancer, genes do not exert their oncogenic or suppressor role in isolation; rather, they are integrated into large, complex, and intertwined regulatory networks. These cancer networks comprise a combination of factors with recurrent gene regulatory structures, termed network motifs, which can cause non-linear dynamics in gene expression levels. This plays a crucial role in determining cellular phenotypes (187–192). SOX10, MITF, and other melanoma development central factors form a myriad of these network motifs, which have relevant implications for tumor progression and therapy resistance. The interaction between SOX10 and MITF is well-documented, establishing a critical link between oncogenes, cellular transcription machinery, and melanogenesis (56). MITF is a pivotal regulator in melanocyte development (193), melanoma progression (194, 195), and immune responses (107), with its activity levels determining melanoma cell phenotypes. Other factors such as TFs (e.g. CREB, and LEF1), miRNAs (e.g. miR-101, miR-137, and miR-218), microenvironmental stimuli (196), and epigenetic states are involved in regulating MITF and SOX10 expression and activity states (105, 196, 197).

Due to the significance of SOX10 and its interacting molecules in melanoma, it is important to analyze it within the broader context of gene regulatory networks. Here, we assembled a gene regulatory network that not only considers the involvement of SOX10 in the control of multipotency (48, 198), but also its involvement in the regulation of glial cell differentiation (48, 199) and melanoma plasticity (51, 193, 200, 201). The network was reconstructed by integrating molecular interactions from gene regulatory networks underlying melanoma (202), pluripotency (203), as well as cells from the same lineage (neural crest) like oligodendrocytes (204) and astrocytes(in-house data). The resulting SOX10-mediated gene regulatory network provides a comprehensive regulon landscape in different cell types and processes that may be relevant to characterize melanoma gene-phenotype mapping (**Figure 2**). To focus the analysis on the role of SOX10 in melanoma, we extracted its direct interactions to reconstruct a small network. We then expanded the network using miRTarBase (version 2022) that provides a repository of experimentally validated miRNA-gene interactions (205) and identified known human TFs (206). We further identified network motifs, such as feedforward loops (FFLs) and feedback loops (FBLs), which comprises SOX10, MITF, and miRNAs. Feedforward loops represent a network motif wherein a regulator can influence the activity or expression of its target via two distinct routes. They have the capacity to suppress noise and induce adaptation in gene expression or activity (189, 207). Feedback loops represent a network motif, whereby a regulator and its downstream targets exert reciprocal effects on each other’s activity or expression. They can give rise to bistability and oscillation in gene expression or activity (189, 207).

**Figure 2.**
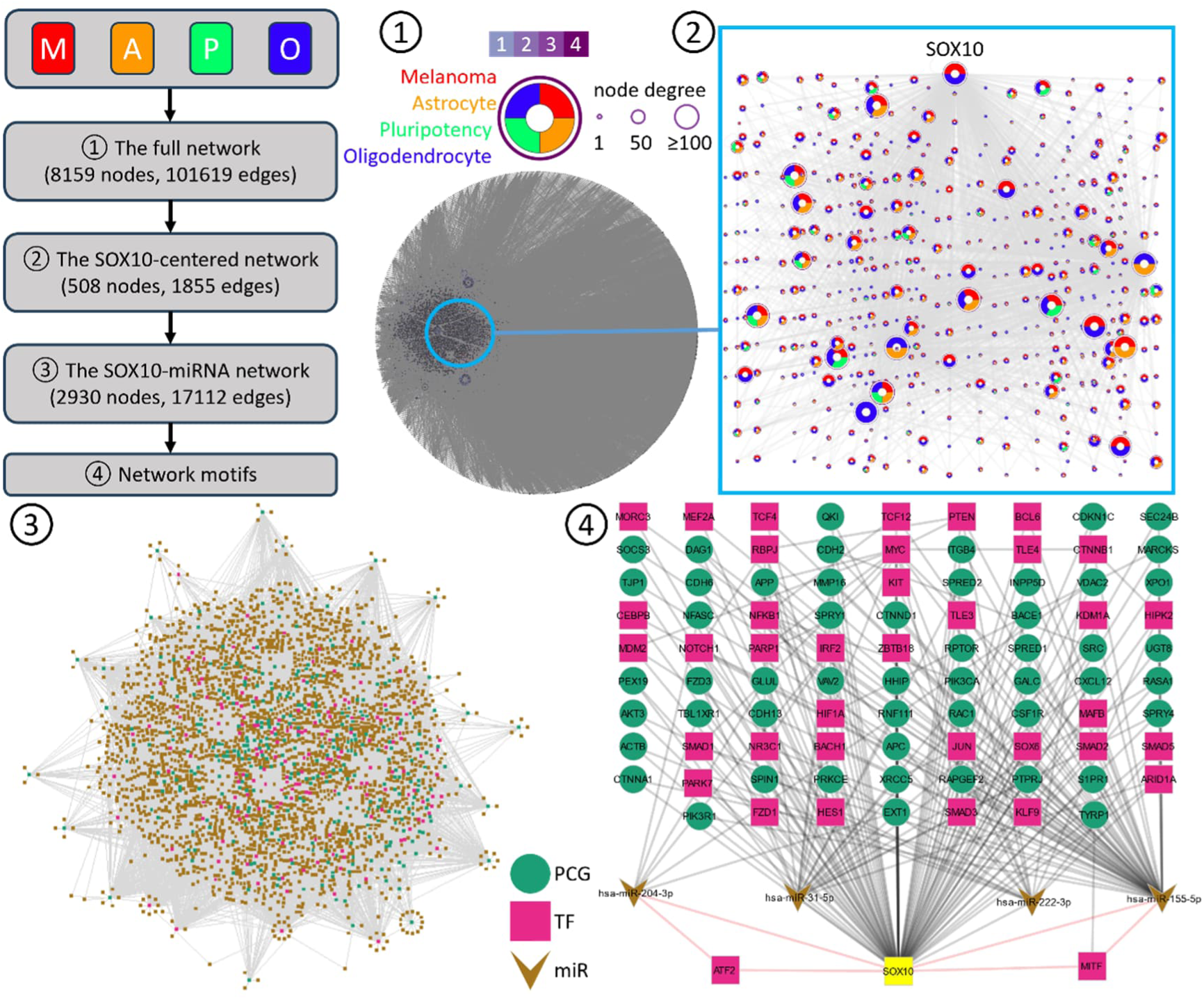
The SOX10-mediated gene regulatory networks and network motifs. The workflow illustrates the four stages of the process used to derive network motifs. The full network was reconstructed in step 1, the SOX10-related interactions were extracted in step 2, miRNA-gene interactions were integrated in step 3, and three-node network motifs were identified in step 4. In the full network and SOX10-centered network, each node is represented by a ring diagram. This diagram indicates the origin networks of the genes associated with that node. These origin networks include melanoma, astrocyte, pluripotency, and oligodendrocyte. Additionally, the node border represents the number of networks to which the gene belongs. The greater the number of networks, the darker the border. The size of a node is proportional to the number of interacting genes, which is referred to as the node degree. In the SOX10-miRNA network and network motif networks, the nodes are color-coded according to their molecular species, with protein-coding genes (PCGs) depicted in green, TFs in red, and miRNAs in brown. SOX10 is highlighted in yellow. The two network motifs (i.e. MIR155-SOX10-MITF and MIR204-SOX10-ATF2) that are discussed in the main text are also highlighted. The Cytoscape file containing all the networks is available in the supplementary materials.

From the network, we obtained feedback and feedforward motifs involving SOX10 and MITF with miRNAs, other TFs, and their mutual targets that were (differentially) expressed in melanoma. As these motifs are regulated by central melanoma gene regulators, we hypothesized that they may play a crucial role in melanoma malignant phenotypes and progression. The following sections present an elaboration on the key gene expression dynamics and their potential for regulating phenotypic plasticity for the most prominent families of network motifs identified during our analysis.

### 3.1 TFs as network hubs in melanoma: The SOX10-mediated MITF gene expression

The SOX10-centered network contained TFs that act as network hubs with high node degrees, making them central to the network’s structure and function (Figure S1 and Table S2). Hub TFs in gene regulatory networks act as master regulators, controlling the expression of numerous genes and thereby driving critical cancer-related processes (208, 209). For example, SP1, HIF-1, and MYC function as master regulators of cancer by regulating essential cancer processes like cell metabolism and proliferation (210, 211). KLF5, a known master regulator across multiple cancer types (212), can influence the tumor microenvironment by regulating T lymphocyte activation and functionality (213). FOXO1 plays a pivotal role in the regulation of mantle cell lymphoma by effectively targeting the cancer’s lineage-specific transcriptional dependencies (214). Additionally, TF hubs interact among themselves, putatively achieving a fine-tuning of the transcriptional programs underlying key phenotypes, which is essential for maintaining cellular homeostasis and responding to oncogenic signals. For example, FOXM1 and CENPF act as synergistic master regulators, promoting tumor growth through coordinated regulation of target gene expression and the activation of key signaling pathways associated with prostate cancer malignancy (215).

In the context of our study, the most pertinent network hubs are SOX10 and MITF, as both TFs play a crucial role in melanoma. Furthermore, SOX10 can directly bind to MITF promoter thereby activating the transcription of MITF (216–218), which, in turn, can modulate a range of genes associated with cancer hallmarks. Theoretically, the gene expression of MITF regulated by SOX10 could be formulated in three models resulting in different dynamics, namely linear, saturated, and sigmoidal (**Figure 3**). Within the framework of the linear model (a.k.a. the rheostat model (195), a linear dependency between the expression levels of MITF and SOX10 is proposed, resulting in a direct and proportional association between their expression levels. Nevertheless, this model may not accurately reflect the reality of MITF regulation by SOX10, as SOX10 has been observed to possess a greater number of binding sites in the promoter of MITF (32). Some of these binding sites demonstrate potential in inducing non-linear dynamics when observed alone or in conjunction with other TFs, such as PAX3 (32). The saturated relationship between SOX10 and MITF expression levels is another potential theoretical framework. In this scenario, as SOX10 levels progressively increase, the corresponding increases in MITF expression levels exhibit a diminishing rate of change for equivalent increments in SOX10. This theory has been widely used to model the dynamics of gene expression regulated by TFs, since an excess of TFs may not have access to their binding sites in the promoter of target genes (219, 220). The sigmoid model shows a sigmoidal relationship between SOX10 and MITF expression. While increments in SOX10 expression induce relatively modest elevations in MITF when SOX10 levels are either exceptionally low or exceptionally high, they elicit substantial alterations in MITF when SOX10 levels reside outside the extreme ranges. TF cooperativity has the potential to induce a sigmoidal shape of transcriptional activation, which is of interest in the context of SOX10, given that this factor has multiple binding sites at the MITF promoter. These binding sites could, therefore, serve to trigger the sigmoidal activation of MITF (221).

**Figure 3.**
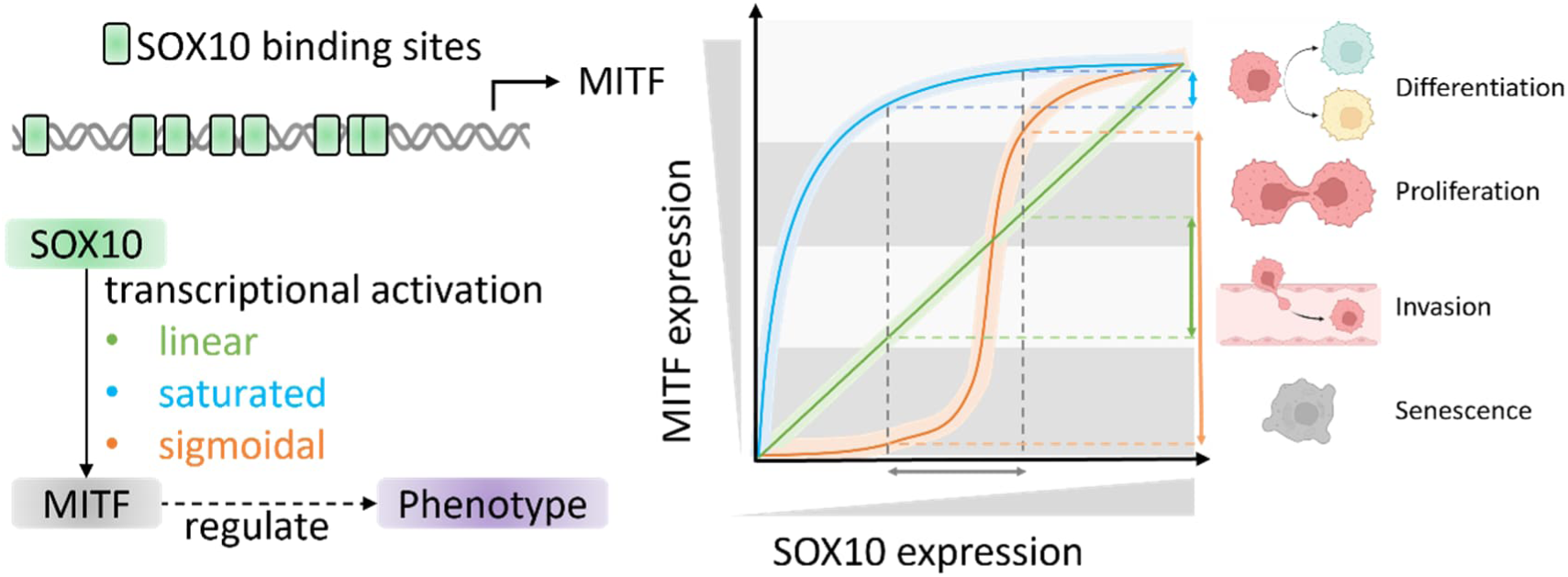
The dynamics of SOX10-mediated MITF expression. The cartoon illustrates the binding sites of SOX10 in the MITF promoter. The diagram illustrates the interaction between SOX10 and MITF, wherein the dynamics of SOX10-mediated MITF expression are proposed in three models: linear (green), saturated (blue), and sigmoidal (orange). The expression levels of MITF affect its downstream genes and determine tumor phenotypes (as elucidated by the rheostat model (195)), including senescence, invasion, proliferation, and differentiation. In the dynamic plot, the solid thin lines represent the average behavior of tumor cells observed in the bulk data, while the shaded thick lines represent the heterogeneous behavior of individual cells observed in the single-cell data. The change in SOX10 expression level (the horizontal grey arrowed line) has different effects on MITF expression levels (the vertical-colored arrowed lines) in different models. The length of the vertical arrowed lines indicates the sensitivity of MITF to the change in SOX10 expression level, which varies depending on the model type and the actual SOX10 expression level. The tumor phenotypes are separated by gray zones, and the zone size may vary for individual cells due to their genomic profiles.

The expression level of MITF is key for determining phenotypes of melanoma cells (195). The heterogeneous expression of MITF in melanoma can be explained by a dynamic process allowing tumor cells to switch back and forth between different phenotypic states (a.k.a. phenotype switch). For instance, studies have shown that tumor cells can switch between migratory and invasive states depending on the progression of tumor cells (195) and anti-tumor treatments can change tumor states (222). The single-cell study investigated tumor heterogeneity in primary melanoma, revealing distinct cell phenotypes associated with high and low expression of MITF (223). Therefore, the proposed relationships between SOX10 and MITF expression levels could explain such biological phenomenon. While the three models (linear, sigmoidal, and saturated) illustrate possible scenarios at the bulk level, they assume identical SOX10 expression within the tumor. However, single-cell data demonstrates diverse SOX10 levels across individual cells (224). Therefore, a more comprehensive explanation for tumor heterogeneity likely involves two key factors: (i) **Differential Sensitivity to SOX10**: Tumor cells exhibit varying sensitivities to SOX10 due to inherent differences in SOX10 expression levels. Heterogeneous expression of SOX10 in individual tumor cells puts them in different states. This can lead to different changes in MITF expression levels for the same increase in SOX10 if the regulation of MITF by SOX10 is non-linear (**Figure 3**). Other TFs promote MITF expression, and the combinatorial cooperation between them in triggering MITF transcription can fine-tune MITF expression and induce single-cell-specific expression. For example, several mechanisms may result in TF cooperation or competition for gene regulation (225, 226). These include direct TF-TF interactions prior to DNA binding and TF-DNA binding that increases the binding affinity of another TF. (ii) **Variations in Genomic Profile-Dependent Models**: It is proposed that individual tumor cells may harbor distinct variations of the proposed models (linear, sigmoidal, or saturated) depending on their unique genomic profiles, which can influence their MITF response to SOX10 (**Figure 3**). This hypothesis is supported by the finding that in seemingly identical cancer cells, dynamic trajectories for a few proteins significantly differ, resulting in different cell fates such as cell death or survival (227). Furthermore, the range of MITF expression levels that define specific tumor phenotypes may also vary between cells. The interplay of these factors contributes to the intricate SOX10-mediated MITF regulation underlying tumor heterogeneity.

### 3.2 Feedforward loops as fine-tuner for regulating gene expression: the MIR155-SOX10-MITF circuit

Gene expression is controlled by a complex network of different regulatory factors, such as TFs regulating gene expression through binding to specific motifs on DNA and miRNAs that regulate gene expression through interacting with the mRNA of genes (228, 229). In this combined transcriptional and post-transcriptional regulation of genes, we identified network motifs such as FFLs in which TFs and miRNAs play a crucial role in regulating the expression levels of genes (230). miRNA-mediated FFLs can be classified coherent and incoherent FFLs, depending on the nature of interactions between their components (231–233). In coherent FFLs, the direct and indirect actions on the targets are consistent, while the two actions are opposite in incoherent ones. In our network, we searched for three node network motifs containing SOX10 and miRNAs (Table S3) and identified a coherent FFL comprising SOX10, MITF, and miR-155 that may play a role in melanoma (**Figure 4**). In this FFL, miR-155 can repress the expression of MITF (234) and SOX10 (235) and SOX10 can upregulate MITF (110). In this section, we elaborated on the dynamic features of the FFL that regulate the expression of SOX10 and MITF.

**Figure 4.**
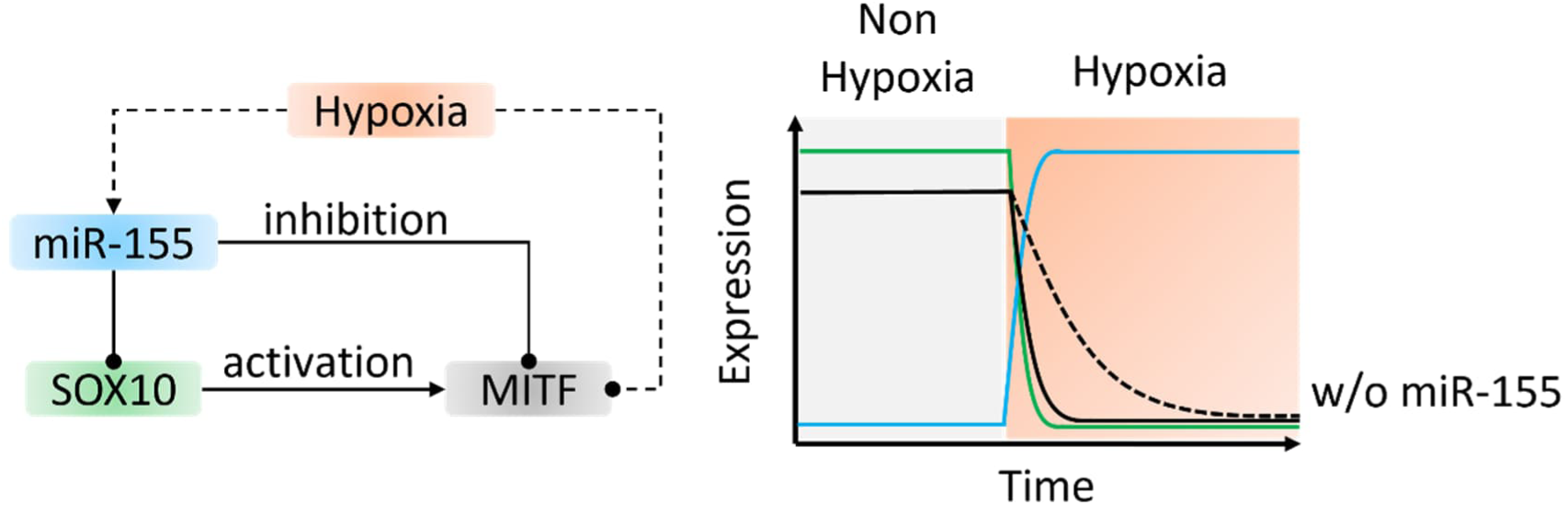
Prevention of transcript leakage regulated by the MIR155-SOX10-MITF FFL. The left figure shows the detailed gene regulation in the MIR155-SOX10-MITF coherent FFL. The oval-head and arrow-head solid lines represent direct inhibition and activation of genes, respectively, while the dashed lines represent indirect gene regulation in hypoxia. The right plot shows the dynamics of FLL gene expression. The occurrence of hypoxia leads to the upregulation of miR-155, which can inhibit both SOX10 and MITF, resulting in the rapid degradation of MITF. In comparison, without miR-155 regulation, the leaky transcripts of MITF will degrade slowly based on its half-life.

#### 3.2.1 Stability in gene expression

It is known that miRNAs can precisely tune the expression levels of their target genes (98, 236, 237). This regulatory function is sophisticated when a miRNA targets a TF and both of them have a common target gene, forming a FFL. The miRNA-mediated FFL not only fine-tunes but also ensures the stability of the target gene expression (238). The dual effects arise from the interplay between transcriptional and post-transcriptional regulation of the target gene by the TF and the miRNA, respectively. For instance, the MIR17-E2F1-RB1 coherent FFL reinforces the stability of RB1’s expression against stochastic fluctuations in E2F1, thus ensuring robust G0/G1 to S transition of cell cycle (238). Most recently, it has been shown that a synthetic miRNA-mediated FFL can stabilize transgenes expression to a desired level or range, making it an attractive option for gene therapies (239). Therefore, in the identified MIR155-SOX10-MITF FFL, miR-155 may stabilize the expression of MITF by facilitating a faster response of MITF to sudden activation or deactivation of SOX10 transcription. This response time of MITF is faster compared to the SOX10-mediated MITF transcription without the involvement of miR-155. This underscores the critical role of miR-155 in stabilizing the expression of MITF via SOX10 to ensure desired phenotypic transition of melanoma cells.

#### 3.2.2 Prevention of leaky gene transcript

The other function of MIR155-SOX10-MITF FFL is to prevent the production of unwanted MITF transcripts (240). miR-155 can repress MITF expression at both the transcriptional level by inhibiting SOX10 and at the post-transcriptional level by binding to MITF mRNA (**Figure 4**). Specifically, low oxygen levels were found to markedly reduce MITF expression in melanoma cells in an indirect HIF-1α-dependent manner, thereby promoting the invasive and metastatic phenotype of tumor cells (241, 242). Meanwhile, hypoxia was found to upregulate the expression of miR-155 (243), which can prevent the translation of leaky MITF transcripts (i.e. already transcribed MITF mRNA in cytoplasm) into proteins. In comparison to the regulation without miR-155, a negative regulator of melanoma cell proliferation and survival (244, 245), the leaky MITF transcripts are still translated into proteins resulting in possible longer retention of tumor cells in aggressive phenotypes that favor tumor progression.

#### 3.2.3 Sign-sensitive delay

In the MIR155-SOX10-MITF FFL, MITF is coherently regulated by miR-155 in two routes – direct and indirect repression (**Figure 5**). Theoretically, the regulation of MITF expression in the FFL can be determined by two types of input function - an AND or an OR gate (231, 232, 246). The AND gate model assumes that the effective repression of MITF can only be achieved when miR-155 reduces the transcription of MITF by SOX10 and concurrently post-transcriptionally represses MITF. This is based on the premise that endogenous upregulation of miRNAs typically exerts mild repressive effects on target genes (247, 248). In contrast, the OR gate model assumes that exogenous upregulation can result in an abundance of miR-155, which can then efficiently repress MITF either by itself or through downregulating its TF SOX10. Regardless of the regulatory route, the repression of MITF remains effective.

**Figure 5.**
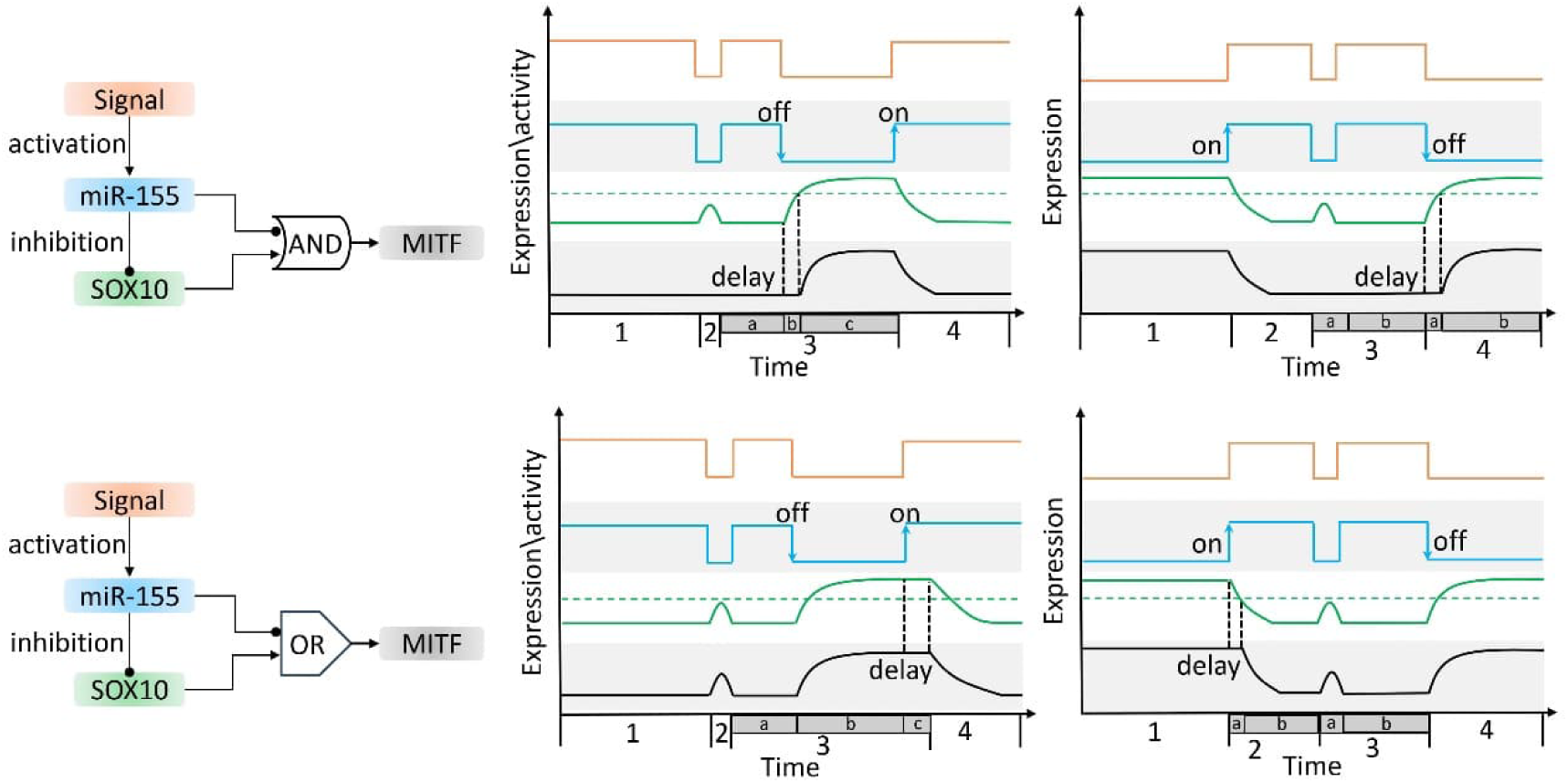
The dynamics of gene expression levels in the MIR155-SOX10-MITF FFL. (**Top**) The coherent FFL with an AND gate function for MITF demonstrates a delay following signal removal (represented by the ‘off’ state at phases 3b and 4a in the middle and right panels, respectively). Furthermore, the FFL functions as a sign-sensitive filter, responding only to persistent signals and not to transient ones (represented by phases 2 and 3a in the middle and right panels, respectively). (**Bottom**) In contrast, the FFL with an OR gate function for MITF exhibits a delay after signal amplification (indicated by the “on” state at phases 3a and 4 in the middle and right panels, respectively) and responds to both transient (phases 2 and 3a in the middle and right panels, respectively) and persistent signals. The colors of the lines correspond to the colors of the components shown in the network motif. The dashed green line indicates the threshold for SOX10 expression level to activate MITF transcription. The numbers on the x-axis are associated with the four phases, which are delineated by segmentations indicated by alphabetic characters. The four gray zones are employed to differentiate the plot areas of the FFL components. To facilitate a more comprehensive understanding of the dynamics of the FFL, a truth table for each plot is provided in the supplementary materials (Figure S2).

The input function of MITF with an AND gate assumes that the effective regulation of MITF by miR-155 requires its direct targeting and indirect regulation via SOX10 (**Figure 5 top left**). This model can cause a delay in the dynamics of MITF expression when miR-155 is turned off by the upstream signal (**Figure 5 top middle**). Initially, miR-155 is highly expressed in response to the upstream signal, while SOX10 and MITF are expressed at low levels due to miR-155’s inhibitory function (phase 1). When the signal is temporarily removed, miR-155 expression is downregulated, resulting in the transient upregulation of SOX10 (phase 2). However, the expression level of MITF remains unchanged because the transient upregulation of SOX10 does not pass the activation threshold required for the transcription of MITF. Therefore, the AND gate ensures stability in MITF expression during a transient downregulation of the upstream signal. When the signal is persistently removed, miR-155 is persistently downregulated, leading to the derepression of MITF by miR-155 (phase 3). However, MITF expression is delayed due to the AND gate function, which requires the expression level of SOX10 to pass the activation threshold first. The length of the delay can be determined by biochemical parameters of SOX10. For instance, the higher the activation threshold for the MITF promoter by SOX10, the longer the delay. When the signal is resumed (phase 4), miR-155 upregulation leads to the silencing of MITF by repressing its expression at both transcriptional levels (via repression of SOX10) and post-transcriptional levels. This results in synchronized dynamics of SOX10 and MITF expressions. This phenomenon is referred to as sign-sensitive delay, which means that the delay occurs depending on the changing direction of the signal (off to on or on to off). In the case of the AND gate model, the delay occurs when the signal is turned off, not when it is turned on. Similar effects can also be expected when the initial expression level of miR-155 is low (**Figure 5 top right bottom;** phase 1). The turning on of the signal results in the upregulation of miR-155, which downregulates SOX10 and MITF (phase 2). The removal of the upstream signal temporarily does not induce a change in MITF expression (phase 3). The delay of MITF occurs when the signal is fully turned off (phase 4). Studies have shown that in biological contexts, TGF-β can modulate miR-155 expression depending on the cell type and tissue environment (187, 249–251). TGF-β is involved in tumor progression and EMT by regulating the MIR155-SOX10 axis (235, 252). The AND gate model of the FFL suggests that a transient change in TGF-β can modulate the expression of miR-155 and SOX10, but MITF is resistant to transient changes in the expression levels of its regulators. Only persistent TGF-β signaling can lead to a change in MITF expression, which occurs with a delay compared to the upregulation of SOX10 expression that can lead to a phenotype switch of tumor cells, for example, from an invasive to a proliferative phenotype (253, 254).

In contrast, the input function of MITF with an OR gate assumes that the modulation of MITF expression necessitates either direct or indirect regulation by miR-155 (**Figure 5 bottom left**). This can result in a different delaying effect in the expression change in MITF. The delay occurs when the signal is turned on not when the signal is turned off (**Figure 5 bottom middle**). Particularly, when the signal persists at a high level at the beginning, miR-155 is highly expressed resulting in the low expressions of SOX10 and MITF (phase 1). When the signal is temporarily turned off, the transient downregulation of miR-155 leads to the upregulation of both SOX10 and MITF (phase 2). In this case, MITF responds promptly to the transient signal change because the OR gate function assumes that the removal of miR-155 repression is sufficient to enable MITF to regain its expression. When miR-155 is persistently downregulated due to the removal of the signal, the expression of MITF is synchronized with the expression of SOX10 (phase 3). When the signal is turned on again, the expression of miR-155 is upregulated, resulting in the downregulation of SOX10 (phase 4). However, once the repressive effect of miR-155 returns, it takes time for the expression level of SOX10 to fall below the activation threshold, thus delaying the decrease in the expression level of MITF. As a result, the response of MITF is delayed when the signal is turned on. Such a delay effect in MITF may also occur when the signal is off at the beginning, then turned on, and finally turned off again (**Figure 5 bottom right**). It is shown that miR-155 is upregulated in response to infection or injury due to various inducing factors, including pathogen-associated molecular patterns and damage-associated molecular patterns (PAMPs/DAMPs) (255), alarmins such as IL-1α (256), and inflammatory stimuli like TNF, IL-1β, and interferons (255), as well as hypoxia (243). In these biological contexts where an upstream signal activates the expression of miR-155, the OR gate model allows for immediate upregulation of MITF via SOX10 when the signal is turned off, inducing a rapid switch to a more malignant tumor phenotype (**Figure 5 bottom middle**). For example, upregulated MITF expression protects melanoma cells from apoptosis, facilitating resistance to treatment (257); high levels of MITF expression activate some genes associated with cell cycle and differentiation, making melanomas more proliferative and differentiated (114). On the other hand, when the signal is turned on (**Figure 5 bottom right**), the delayed response in MITF expression can maintain the tumor in the malignant phenotype for a longer time, favoring tumor progression.

### 3.3 Feedback loops underlying melanoma aggressiveness: the SOX10-MIR204-ATF2 circuit

In addition to FFL, feedback loops (FBL) is another type of network motifs that exist in molecular networks such as gene regulation and signal transduction networks underlying tumor genesis and tumor progression (258). FBLs are regulatory motifs, in which a downstream molecule in a regulatory circuit can promote (positive) or inhibit (negative) an upstream process. Negative FBLs are responsible for maintaining homeostasis, which is the ability of gene circuits to balance noise (189, 207) and the ability to promote quick cessation to avoid overwhelming circuit activation. Positive FBLs can induce signal amplification or, when combined with mutual gene repression, elicit switch-like behaviors in gene activation. By analyzing our network, we searched for FBLs containing SOX10 and miRNAs (Table S3) and identified a FBL comprising of miR-204, ATF2, and SOX10 that can regulate the expression of MITF. In the FBL, miR-204 can post-transcriptionally inhibit the expression of ATF2 observed in glioblastoma cells (259) and ATF2 can inhibit SOX10 expression in melanoma cells (260), while SOX10 can upregulate the transcription of miR-204 as reported in oligodendrocytes (259). In a mouse model of melanoma, Scah et al. demonstrated that ATF2 negatively regulates MITF transcription through downregulation of SOX10 (260). Downregulation of ATF2 can upregulate the expression of MITF via SOX10 and transform melanoma into a metastatic phenotype. Except for ATF2, the expression of SOX10 can be modulated by other signals and factors, such as phosphorylation of SOX9 by AKT (261), deletion of SLK (262), MAPK signaling pathway (263), or recurrent open chromatin domain (264).

From a dynamic perspective, the SOX10-MIR204-ATF2 positive FBL can function as a toggle switch, resulting in bistability in gene expression levels and regulating the switch between two steady states that are related to different tumor phenotypes. It has been recently reported that melanoma can show three cell states - melanocytic and mesenchymal cell states and an intermediate cell state distinct from the other two (154). Such kinds of intermediate cell states are also identified EMT, which was regarded as a binary process but showed distinct intermediate states in pancreatic and epidermoid carcinoma (265, 266). Here, we explained how the FBL could regulate the phenotypic switch between the three melanoma cell states (**Figure 6**).

**Figure 6.**
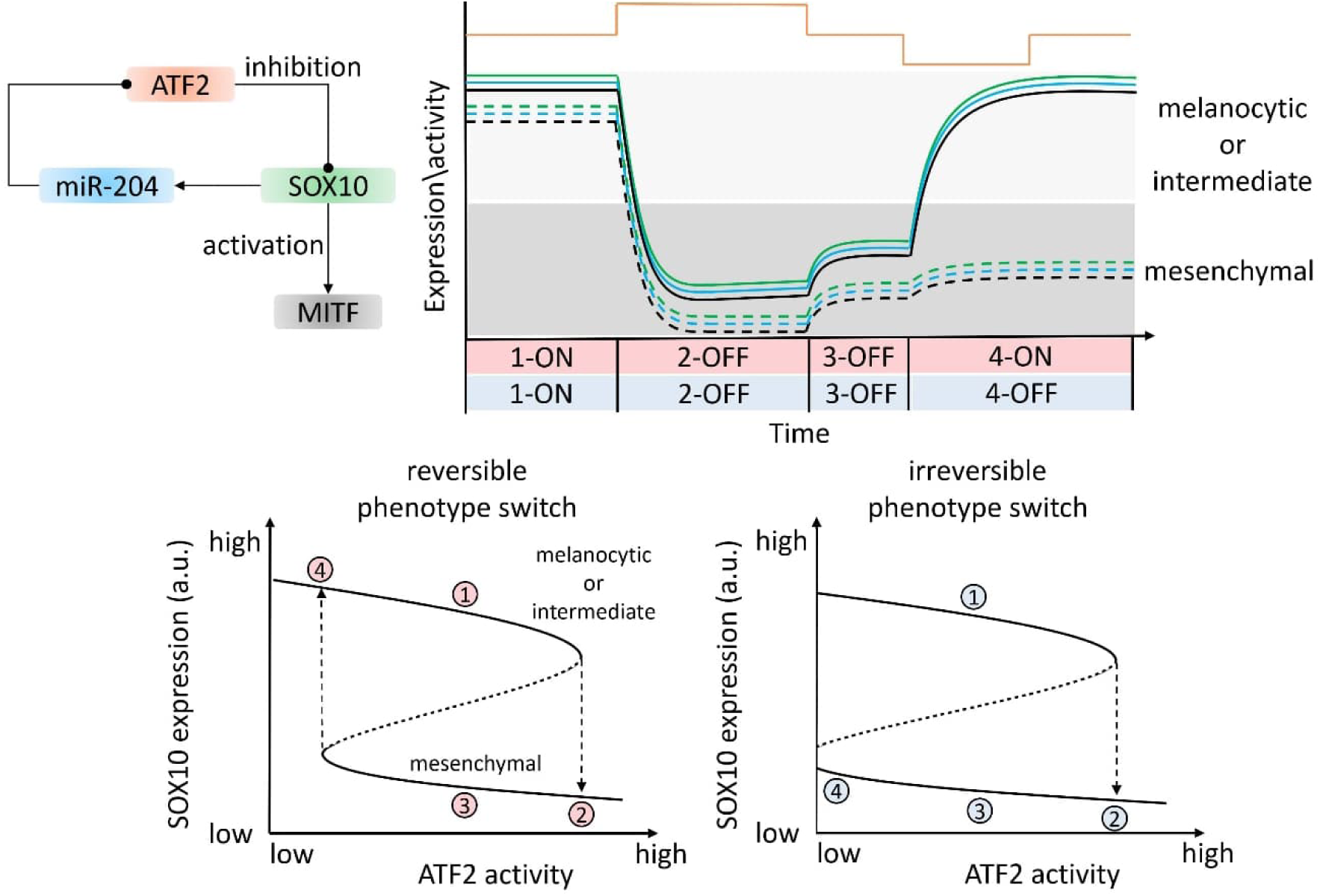
The bistability of gene expression in the MIR204-SOX10-ATF2 FBL. (**Top**) The diagram shows possible molecular interactions in the SOX10-miRNA FBL. The dynamic plot shows the change in activity or gene expression levels in the FBL over time. For simplicity, we assumed that the dynamics of miR-204 and MITF are like SOX10 because SOX10 positively regulates their transcription. The activity of ATF2 is shown at the top of the plot. The colors of the lines correspond to the color of genes shown in the network motif. The solid and dashed lines correspond to the scenarios shown in bottom left and right panels, respectively. The background colors indicate phenotypic states of melanoma cells that is determined by the expression level of MITF. The numbers on the x-axis correspond to the four phases described in the main text and they also correspond to the numbers in the bottom plot. The terms ‘ON’ and ‘OFF’ indicate the upper and lower steady states of SOX10, respectively. (**Bottom**) The bifurcation plot illustrates the reversible (left panel) and irreversible (right panel) switching between melanoma cell states. The solid and dashed parts of the curves represent stable and unstable steady states of SOX10, respectively. The upper and lower steady states correspond to ON and OFF steady states in the middle plot. The numbers in red and blue circles indicate the dynamics of gene expression and follows the order as shown in the middle plot (i.e. 1->2->3->4). The ability of melanomas to revert to their initial cell states is dependent on the location of the left bifurcation point. If the left bifurcation point is located within the activity interval of ATF2, the tumor cells can switch back to their initial states. If the left bifurcation point is in a position that lacks biological meaning (e.g., a negative value of ATF2 activity), melanomas become trapped at the lower part of the curve and cannot return to their initial states. For a more detailed theoretical explanation about bifurcation points, please refer to (278).

When ATF2’s activity is low, SOX10 is expressed at a high level (phase 1-ON steady state) resulting in high expression of miR-204 and MITF that make melanoma cells show a melanocytic or an intermediate phenotype. When the activity of ATF2 increases, the expression of SOX10 is downregulated (phase 2). This, in turn, downregulates the expression of miR-204 amplifying the activity of ATF2 due to the positive FBL. The decreased expression level causes SOX10 to switch to a lower steady state (phase 2-OFF steady state), resulting in downregulation of MITF. The low expression level of SOX10 and MITF causes the tumor to show a mesenchymal phenotype (260, 267). When the SOX10 expression level stays at the OFF steady state, it cannot revert to the ON steady state even when the ATF2 activity returns to the low level (phase 3-OFF steady state). Consequently, the SOX10 expression level remains relatively low, leading to a relatively low MITF expression that causes the tumor to continue exhibiting a mesenchymal phenotype. Finally, the expression of SOX10 can only return to the upper steady state when the ATF2 activity continues decreasing and crosses below its initial level (phase 4-ON). The resumed expression level of SOX10 leads to the upregulation of miR-204, which can further repress ATF2, resulting in normal expression of MITF that may revert tumor cells to a melanocytic phenotype (**Figure 6 bottom left**). This phenomenon is called hysteresis (268), in which the phenotypic switching of melanomas lags behind changes in ATF2 activity. However, it remains unknown whether the transition of melanoma cells from mesenchymal states towards intermediate or melanocytic cell states is possible(269). Thus, it is also possible that melanoma cells are confined to the mesenchymal phenotype because of the irreversible switching to the melanocytic or intermediate state (**Figure 6 bottom right**). A more complicated scenario could be that several FLBs and FFLs mediated by SOX10, and miRNAs are coupled together to cause tristability in SOX10 expression, resulting in a complex transition among the three melanoma cell states. Such a phenomenon has been observed in EMT and analyzed using mathematical models (270–272). Furthermore, there is ongoing debate regarding the existence of melanoma cell states. Some propose that melanoma cells exist in a spectrum of states in which cell states are continuously changing (273), while others believe that distinct melanoma phenotypes can be stabilized by FBLs (274). Our results align with the latter hypothesis, as the bistability of MITF resulted from the FBL may sustain specific cell states. This phenomenon has also been observed in a FBL consisting of MITF, BRN2 and miR-211, which can result in populations with distinct MITF and BRN2 profiles (275).

Taken together, we showed a potential FBL containing the interactions between SOX10 and miRNAs and elucidated on its role in regulating the dynamics of gene expression. This suggests that the distinct melanoma cell states are underpinned by SOX10-mediated network motifs.

## Discussion and conclusion

### Heterogeneous SOX10 expression levels and complex role of SOX10 in melanoma

The expression patterns of MITF and SOX10 are critical in melanoma progression. However, the heterogeneity of their expression levels within melanoma tumors can significantly impact the dynamics of network motifs. A recent study has demonstrated that individual primary melanoma cells exhibit varying expression profiles for SOX10 and MITF (224), indicating gene expression variability of both TFs that may result in distinct regulatory states. These states can affect downstream signaling pathways and cellular behaviors. Understanding this heterogeneity and its effects is crucial for comprehending gene regulatory programs at the single-cell level and designing cell-based therapies for melanoma. Furthermore, it is important to be aware of the multifaceted and complex role of SOX10 in melanoma. SOX10 can promote the proliferation and survival of melanoma cells, but it can also induce a more differentiated and less aggressive phenotype in these cells (51, 54, 263). A recent study showed that depletion of SOX10 can sensitize melanoma cells to T cell-mediated killing and induction of cell death by cytokines such as TNF-α or IFN-γ, suggesting its ability to regulate the interaction between immune cells and melanoma cells (254). Therefore, additional efforts are required to clarify the function of SOX10-miRNA interactions in melanoma, given the contradictory role of SOX10 and its complicated regulation by miRNAs.

### Interplay between SOX10 and non-coding RNAs

We showed the indispensable role of miRNA in regulating the expression of SOX10 in melanoma. However, the complexity of miRNA-mediated gene regulation is often underestimated, which complicates the underlying molecular mechanisms (276). For example, miRNAs can have off-target effects due to their unspecific binding to target mRNAs, potentially leading to unspecific gene regulation (277). Off-target effects can lead to undesired or unknown effects in melanoma when miRNAs are aberrantly expressed. Therefore, a strategy that achieves the same or greater repressive effect on the target gene while requiring fewer miRNA doses can reduce the unwanted effects caused by miRNA off-targeting. One possible solution is to use cooperative miRNA pairs that can repress a target gene to a desired expression level (96). This approach only requires moderate overexpression of the cooperative miRNAs, compared to the overexpression of a single miRNA. Additionally, it is important to recognize that other ncRNAs can also interact with SOX10. Long ncRNAs, circular RNAs, and other emerging classes of ncRNAs have been shown to be critical in the development and maintenance of cutaneous homeostasis and function, and their dysregulation can contribute to cutaneous malignancies (178). This is important because their involvement in SOX10-mediated gene regulatory networks can complicate molecular regulons in melanoma. Hence, future research focusing on the crosstalk between SOX10 and diverse ncRNAs will provide us with a more comprehensive picture of melanoma pathogenesis and progression.

### The interplay of SOX10 and miRNAs in melanoma

In this paper, we investigated the intricate relationship between SOX10 and miRNAs and elucidate their pivotal role in the pathogenesis of melanoma. We further explored the dynamic interplay that influences the expression of SOX10 and MITF, both of which are integral to melanoma progression. This is achieved through the lens of miRNA-mediated network motifs, specifically FFLs and FBLs, which offer a comprehensive view of the regulatory mechanisms involved.

By integrating network biology methods with dynamical systems theory, we demonstrated the effectiveness of this approach in interpreting experimental findings reported in the literature. This integrative approach provides a quantitative and precise understanding of the regulatory mechanisms that underlie tumorigenesis and tumor progression in melanoma. We identified the driving network motifs that regulate the expression of SOX10 and miRNAs, and their targets. Quantitative analysis is employed to elucidate how the genes involved in the identified network motifs influence the phenotypic plasticity of melanoma.

In conclusion, we highlighted the significance of a network biology perspective in comprehending the complex regulatory mechanisms in melanoma, which lead to better understanding of SOX10 regulation by miRNAs in melanoma. SOX10 and miRNAs could be potential therapeutic targets for melanoma due to their widespread expression in tumor samples and their instrumental role in modulating the hallmarks of cancer.

## Acknowledgements

This work has been supported by the German Ministry of Education and Research (BMBF) through the initiatives e:Med-MelAutim on cancer and autoimmunity [01ZX1905A to XL and JV] and KI-VesD on computation modelling and artificial intelligence guided cancer diagnostics [031L0244A to JV]. XL also acknowledges the support from the Johannes und Frieda Marohn-Stiftung [Alz/Iko-Lai/2022]. In addition, we thank the open access support funding from Tampere University, Finland.

## Author Contribution

Conceptualization: XL; Data curation: XL, CYL; Formal analysis: XL; Funding acquisition: XL, JV; Investigation: XL; Methodology: XL; Project administration: XL; Resources: XL, CB, JV; Software: XL, CYL; Supervision: XL, JV; Validation: XL; Visualization: XL, CYL, ZSZ; Writing-original draft: XL, CYL; Writing-review & editing: XL, CYL, AW, MH, CB, JV.

## Data Availability Statement

The supplementary materials and code for this work will be deposited at Zenodo (the Zenodo doi will be provided after the paper is accepted for publication) for reproducing the results and promoting the reuse of the computational workflow. The following supplementary files are provided:

1. Figure S1 shows the Boolean truth table underlying the gene expression dynamics in Figure 5.
2. Figure S2 shows the node degree distribution of transcription factors and protein-coding genes within the SOX10-centered network.
3. Table S1 contains a comprehensive list of references utilized for the annotation of molecular interactions depicted in Figure 1.
4. Table S2 presents the node degree of transcription factors and protein-coding genes for drawing Figure S1.
5. Table S3 contains the identified three-node network motifs comprising SOX10 and miRNAs.

## Supplementary figures

**Figure S1.**
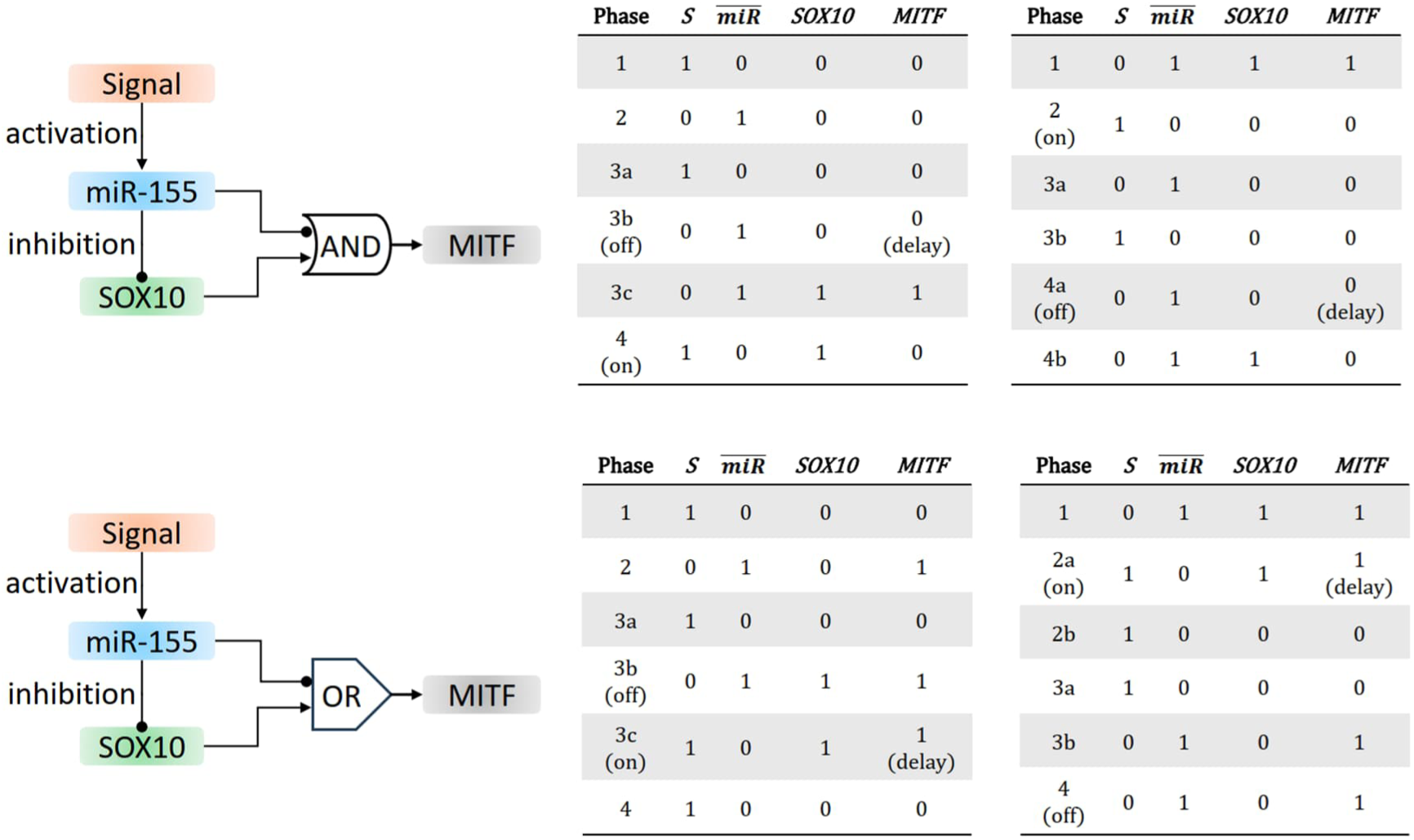
The Boolean truth table of the AND and OR gate model for the MIR155-SOX10-MITF FFL. Each table corresponds to a dynamic plot shown in Figure 2. The phase column represents the four phases, with “on” and “off” corresponding to the states in the dynamic plot. The “delay” corresponds to the observed delay in MITF expression dynamics. 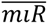 represents the logic NOT expression of miR-155 (i.e. 0->1 or 1->0) because the miRNA inhibits the expression of MITF.

**Figure S2.**
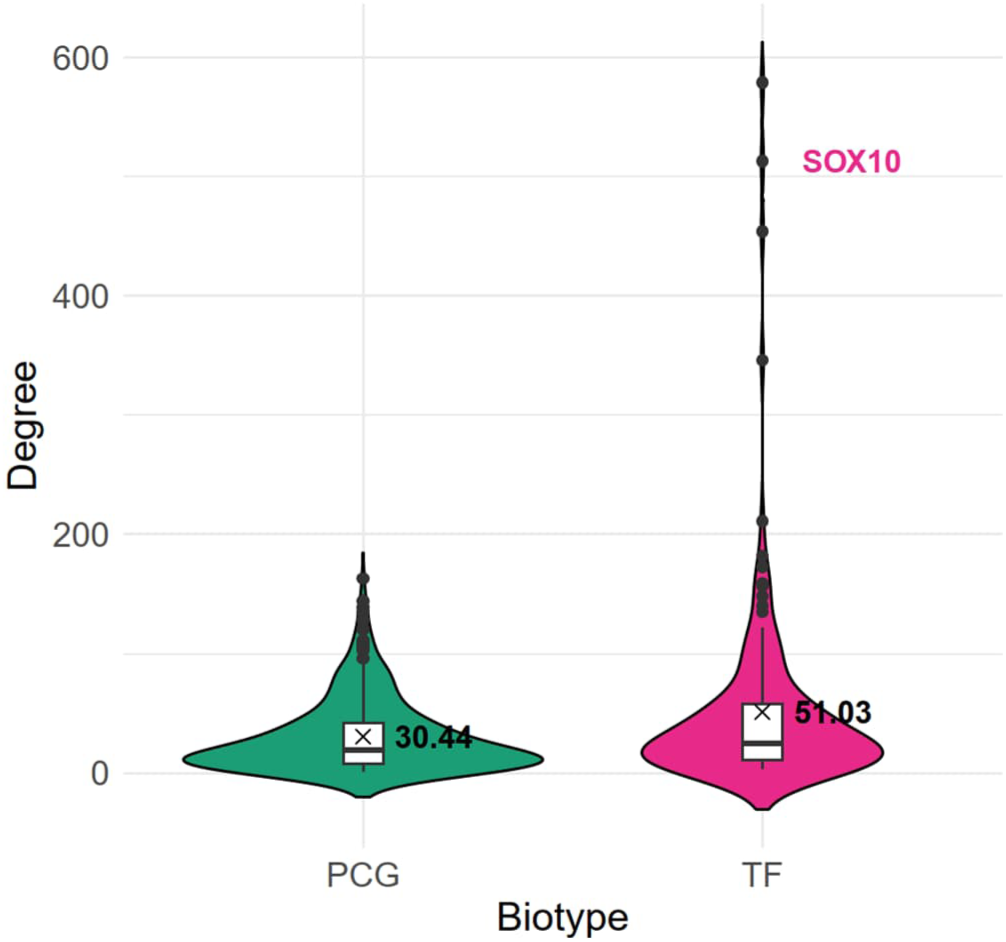
The node degree distribution. The violin plot depicts the distribution of node degree among protein-coding genes (PCG) and transcription factors (TF) within the SOX10-centered network. The embedded boxplot illustrates the quartile distribution of the data, with thick horizontal lines representing the median value and crosses representing the mean value, for which the actual value is provided. The dots represent outliers, with SOX10 highlighted. The TFs have higher node degrees than the PCGs.

## Supplementary tables

**Table S1** presents a comprehensive list of references utilized for the annotation of the molecular interactions depicted in Figure 1. Reaction_ID: This is the unique identifier for a given reaction. It can be enabled by navigating to “View” and then selecting “Show reaction ID” in CellDesigner. It should be noted that the reaction ID is not continuous, as some edges were deleted (i.e. re20, re70, and re74) when the map was created. Ref1-5: These columns display the full bibliographic information of an article that was used to annotate the edge in question. It is possible for a single edge to contain multiple references.

**Table S2** presents the node degree of transcription factors (TFs) and protein-coding genes (PCGs) for drawing Figure S1. SOX10 is highlighted.

**Table S3** contains the identified three-node network motifs comprising SOX10 and miRNAs. The two motifs discussed in the main text are highlighted.

